# The Arabidopsis *INNER NO OUTER (INO)* gene acts exclusively and quantitatively in regulation of ovule outer integument development

**DOI:** 10.1101/2022.08.12.503699

**Authors:** Debra J. Skinner, Trang Dang, Charles S. Gasser

## Abstract

The *INNER NO OUTER* (*INO*) gene is essential for formation of the outer integument of ovules in *Arabidopsis thaliana*. Initially described lesions in *INO* were missense mutations resulting in aberrant mRNA splicing. To determine the null mutant phenotype we used CRISPR to induce frame-shift mutations and found, in confirmation of results on another recently identified frame-shift mutation (Vijayan et al., 2021), that such mutants have a phenotype identical to the most severe splicing mutant (*ino-1*), with effects specific to outer integument development. We show that the altered protein of an *ino* mRNA splicing mutant with a less severe phenotype (*ino-4*) does not have INO activity, and the mutant is partial because it produces a small amount of correctly spliced *INO* mRNA. Screening for suppressors of *ino-4* in a fast-neutron mutagenized population identified a genetic duplication of the *ino-4* gene leading to an increase in the amount of this mRNA. The increased expression led to a decrease in the severity of the mutant effects, indicating that the amount of INO activity quantitatively regulates outer integument growth. The results further confirm that the role of *INO* in Arabidopsis development is specific to the outer integument of ovules where it quantitatively affects the growth of this structure.

## INTRODUCTION

As the structures that develop into seeds, ovules are an essential component of sexual reproduction in seed plants. Angiosperm ovules comprise a combination of diploid maternal tissues and the haploid megagametophyte that develops to form the embryo and endosperm following double fertilization. Most angiosperm ovules include two sheathing maternal structures, the integuments, which appear to be essential for aspects of fertilization and/or embryo development and that further develop themselves, forming the seed coat. Genes involved in regulation of integument formation have been described in *Arabidopsis thaliana* through mutant and molecular studies (reviewed in (Gasser and Skinner, 2019)). First described in Arabidopsis, the *INNER NO OUTER* (*INO*) gene is essential for normal outer integument development, and encodes a protein in the YABBY gene family (Baker et al., 1997; Villanueva et al., 1999). Only two different lesions in the Arabidopsis *INO* locus were identified in extensive forward mutant screens. Both are single base substitutions in introns that result in new upstream splice acceptor sites. While being independently isolated, the *ino-1* and *ino-2* alleles have identical substitutions in intron five creating an AG doublet that when utilized as a splicing acceptor site adds eleven bases to the mRNA leading to a frame-shift near the start of the essential YABBY domain (Villanueva et al., 1999). This *ino* lesion leads to a complete absence of the outer integument in homozygous mutants and near complete female infertility. Another *ino* allele, *ino-4*, has a substitution in intron four, also introducing an AG doublet that acts as a splice acceptor site. In this case, fifteen bases are inserted into the mRNA resulting in addition of five amino acids in the YABBY domain of potential translation product, while still maintaining the reading frame for the remainder of the protein (Villanueva et al., 1999). *ino-4* has a less severe phenotype than *ino-1*, exhibiting incomplete growth of the outer integument and production of approximately one fourth as many seeds as wild-type (Villanueva et al., 1999). This was hypothesized to be due to partial activity of the protein with inserted amino acids (Villanueva et al., 1999).

The altered splicing nature of the only two *ino* mutations initially described in Arabidopsis raised several questions. Both described mutations resulted in introduction of splice acceptor sites that produce aberrant mRNAs. However, both also retain the original splice acceptors raising the possibility that both could still produce wild-type *INO* mRNA. The assay used to indicate that normal *INO* mRNA was not produced, direct sequencing of synthesized cDNA (Villanueva et al., 1999), is not especially sensitive and low levels of wild-type mRNA could have been present. Thus, the actual *ino* null mutant phenotype in Arabidopsis was not clear. It is possible that such a mutation could produce deleterious pleiotropic effects precluding isolation. To assess this we set out to identify null mutant alleles and used CRISPR-mediated mutagenesis to create null mutants. Prior to completion of this manuscript, another group used the same method to identify a putative null mutant and showed that it did not have such pleiotropic effects (Vijayan et al., 2021). Another question arising from the nature of the initially identified mutants is whether the *ino-4* weak mutant effect results from a protein with reduced activity, or from partial leakage of wild-type mRNA formation in this mutant. Finally, we do not know if levels of INO activity differentially effect outer integument formation.

To address these questions, we have re-examined the existing *ino* mutants by additional means, have obtained and characterized additional mutants that include likely null mutants, and have isolated a second site mutant that addresses the question of quantitative expression of *INO*. As also shown in a recent study (Vijayan et al., 2021), we find that newly isolated null mutants duplicate the phenotypic effects of the *ino-1* mutant, indicating that *ino-1* is also effectively a null mutant. We show that the altered protein of *ino-4* does not have INO activity, and the phenotype is weak because it produces a small amount of normal *INO* mRNA. A gene duplication leading to a quantitative increase in the amount of this mRNA leads to a decrease in the severity of the effects, indicating that the amount of INO activity quantitatively regulates the developmental effects of INO. The results further confirm that the role of *INO* in Arabidopsis development is specific to the outer integument of ovules.

## MATERIALS AND METHODS

### Plant material

*Arabidopsis* plants were grown under long day conditions as previously described (McAbee et al., 2006). Pre-existing alleles used in this study are *ino-1* (in the Ler background) and *ino-4* (in the Col-0 background) (Villanueva et al., 1999), and *ino-9* (SALK_0063818) (Alonso et al., 2003). Plant transformation utilized *Agrobacterium* GV3101 (Koncz and Schell, 1986) and the floral dip method (Clough and Bent, 1998).

### Plasmid Manipulation

*pJC02 -* To clone the cDNA for the aberrantly spliced *ino-4* mRNA, cDNA was synthesized from *ino-4* mRNA using Superscript II reverse transcriptase and oligo dT primers (ThermoFisher, Waltham, MA). The specific cDNA was amplified by PCR using Phusion DNA polymerase (New England Biolabs, Ipswich, MA) and primers AtINOcDNA-f and AtINOcDNA-r (Supplemental Table S1), which introduce flanking BamHI and XbaI sites, respectively. The resulting PCR product was inserted into pJet2.1 using the CloneJet cloning kit (ThermoFisher, Waltham, MA) and one sequence-verified *ino-4*-specific cDNA clone was designated pSR06. The cDNA in pSR06 was transferred into pJRM42 (a derivative of pRJM33 – the expression vector for the wild-type INO cDNA from the wild-type INO promoter (Meister et al., 2005)) as a BamH1/Xba1 fragment at these same sites creating pJC01. pJC01 is identical to pRJM33 except having the ino-4 cDNA replacing the wild-type INO cDNA. The NotI insert fragment of pJC01 containing the complete promoter cDNA expression cassette was inserted into the NotI site of pMLBART (Gleave, 1992) to create the plant transformation vector pJC02 which can be selected in plants for phosphinothricine resistance.

*CRISPR mutagenesis* - For CRISPR-mediated mutagenesis we utilized the dual guide RNA approach of Pauwels et. al (2018). Two guide RNA targeting sequences are inserted into two similar vectors at *Bpi*I sites where they are adjacent to Arabidopsis U6 RNA promoters and other constant sequences of the guide RNA, and these two transcription units are subsequently inserted into a plant transformation vector by site-directed recombination using LR Clonase II (ThermoFisher, Waltham, MA) (Pauwels et al., 2018). In our case the synthetic guide RNA targeting sequences INOCRIS4 and INOCRIS5 (Supplemental Table S1) were inserted into the *Bpi*I sites of pMR218 (Pauwels et al., 2018) and a derivative of pMR217 (Pauwels et al., 2018) with a synthetic U6 RNA promoter. LR Clonase II was used to insert the transcription units from these two plasmids into a derivative of plant transformation plasmid pDE-CAS9Km (Pauwels et al., 2018) in which the kanamycin resistance gene is replaced by a gene expressing phosphinothricine acetyl transferase (conferring resistance to BASTA herbicide), and also incorporating a gene for production of the green fluorescent protein (eGFP) in the embryo under control of the Arabidopsis oleosin promoter (Plant et al., 1994), resulting in pGD60. GFP fluorescence in the embryo from the P-Oleosin::eGFP gene allows for the differentiation of seeds harboring the CRISPR plasmid from those which have lost the plasmid through genetic segregation (Aliaga-Franco et al., 2019).

### Polymerase Chain Reactions

Polymerase chain reactions utilized DreamTaq polymerase (ThermoFisher, Waltham, MA) and the supplied reaction buffer components. Superscript II reverse transcriptase (RT) (ThermoFisher, Waltham, MA) with oligo d(T) primers was used for cDNA synthesis for reverse transcriptase (RT)-PCR.

### Microscopy

Scanning electron microscopy was performed as described (Kelley et al., 2009). Dark field images were acquired on a Zeiss Axioplan microscope (Carl Zeiss Inc, Oberkochen, Germany) with a Moticam 5 digital camera (Motic Inc., San Antonio, TX). Images were cropped and matched for contrast in Adobe Photoshop CS5 (Adobe, San Jose, CA).

### Fast Neutron Mutagenesis and Mutant Identification

Homozygous *ino-4* seeds were mutagenized with fast neutrons (60 Gy) at the University of California, Davis McClellen Nuclear Research Center, Sacramento CA. The treatment resulted in 20 – 30% lethality and 10 – 20% of the surviving seed produced fully infertile plants. M1 seeds were planted in pools of five plants. M2 seed from each pool were collected and fifty seeds from each pool were planted, and grown to anthesis. Ovules were examined under a stereomicroscope to detect deviations from the *ino-4* mutant phenotype.

### Genetic Mapping and Whole Genome Analysis

To map the location of the mutation responsible for the suppressed *ino-*4 phenotype (*i20*), we outcrossed the *i20/i20 ino-4/ino-4* plants to L*er*. F2 progeny from this cross were used to map the location of *i20* to Chromosome 3 using Cleaved Amplified Polymorphic Sequences (CAPS), and Simple Sequence Length Polymorphism (SSLP) markers (Jander et al., 2002) and indel markers (Hou et al., 2010). The suppressed phenotype observed in the *i20* mutation acted semi-dominantly, thus individuals used for fine mapping were progeny tested, and only homozygous individuals were used for further analysis. The mapped location of *i20* was between marker nga126 (3.098 Mb) and 3-4608 (4.608 Mb). Twelve individuals from F2 and F3 mapping populations, homozygous for *ino-4* and *i20*, were used to extract DNA for sequencing.

Leaf samples from *ino-4* and *ino-4 i20* plants were separately pooled and DNA extracted with Nucleon Phytopure DNA extraction kit (GE Healthcare, Pittsburgh, PA), according to the provider’s instructions. NEBNext DNA library components (New England Biolabs, Ipswich, MA) were used for fragmentation to 200-500 bp, and library preparation with Nextflex-96 indexes (Bioo Scientific, Austin, TX).

Libraries were pooled and sequenced using the PE100 bp protocol on a single lane of Hiseq 2000 (Illumina, Inc., San Diego, CA) at the Vincent J. Coates Genomics Sequencing Laboratory at UC Berkeley. 67.5 and 99.9 million reads were obtained for *ino-4* and *ino-4 i20* respectively. Reads were preprocessed to remove tag and poor-quality regions using Sickle (Version 1.33, https://github.com/najoshi/sickle), and Scythe (https://github.com/najoshi/scythe), respectively. Reads were aligned with the Col-0 TAIR 10 reference sequence (Lamesch et al., 2012) using BWA-MEM (Li, 2013). Aligned sequences were viewed in Tablet (Milne et al., 2013) to visualize changes relative to the wild-type Col-0 sequence in the region identified as including the *i20* suppressor based on the mapping.

### Data availability

Newly derived plant lines have been deposited at the Arabidopsis Biological Resource Center at Ohio State and other plant materials are available from the cited sources from which they were obtained. Raw genomic sequence data has been deposited in the NCBI SRA database under accession PRJNA866995.

## RESULTS

### *ino-4* mis-spliced mRNA cannot support outer integument growth

While the *ino-1* mutant exhibits no growth of the outer integument, the *ino-4* mutant shows partial growth of the outer integument ((Villanueva et al., 1999), Fig. 1B, 1C, Fig. 2B, 2C). This was hypothesized to result from partial INO activity of the protein product of the mis-spliced *ino-4* mRNA. This product has a fifteen-nucleotide insertion that would result in insertion of five amino acids at Glu-147 in the translated protein and no other change in the reading frame (Villanueva et al., 1999). To determine whether the mutant protein has activity, we tested transgenic complementation of *ino-1* with a construct that produces the *ino-4* mutant protein. The wild-type cDNA was shown to be effective at complementing *ino-1*. A construct, pRJM33 (Villanueva et al., 1999), including 2.3 kb of *INO* 5’-flanking DNA, the *INO* cDNA, and 2.0 kb of *INO* 3’-flanking DNA led to formation of fully wild-type ovules in twenty-eight of forty-two independently produced transgenic homozygous *ino-1* plants (Meister et al., 2005). When the wild-type *INO* cDNA was replaced in pRJM33 with the aberrantly spliced cDNA of *ino-4*, and the resulting construct (pJC02) was introduced into plants, twenty-nine of twenty-nine independently transformed homozygous *ino-1* transgenics retained the *ino-1* ovule phenotype. These results differ significantly (P < 0.0001, two-tailed Chi^2^) demonstrating the failure of the *ino-4* cDNA sequence to complement the mutant, and hence the failure of the extended ino-4 protein to function.

**Fig. 1.**
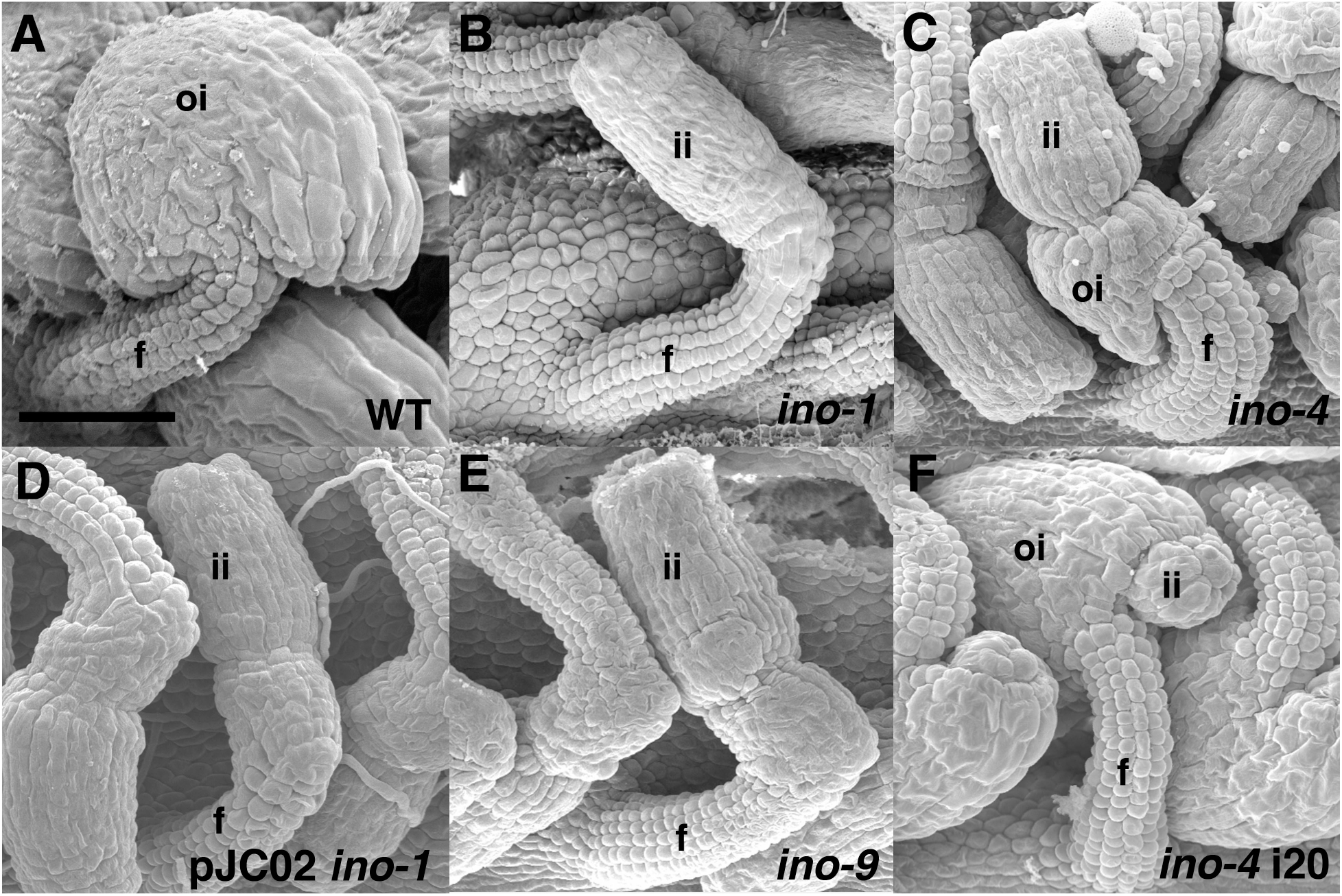
Scanning electron micrographs of Arabidopsis ovules. All ovules are from flowers at or just following anthesis and are oriented with the gynoapical side to the right. A) Wild-type ovule where the curved outer integument has covered the inner integument. B) *ino-1* mutant ovule where the outer integument is absent, exposing the inner integument which remains erect due to this absence. C) *ino-4* mutant ovule has a rudimentary outer integument structure which fails to cover the inner integument. D) ovules of *ino-1* mutant plant containing transgene transcribing the mutant *ino-4* cDNA (pJC02) that fails to complement so that the ovules retain the *ino-1* phenotype. E) *ino-9* mutant ovules that are phenotypically indistinguishable from those of *ino-1*. F) i20 partially restores outer integument growth of *ino-4* so that in the *ino-4* i20 double mutant the outer integument can largely cover the inner integument, leaving only a small part of the inner integument protruding. f, funiculus; ii, inner integument; oi, outer integument. Bar in A) is 50 µm in all panels.

**Fig. 2.**
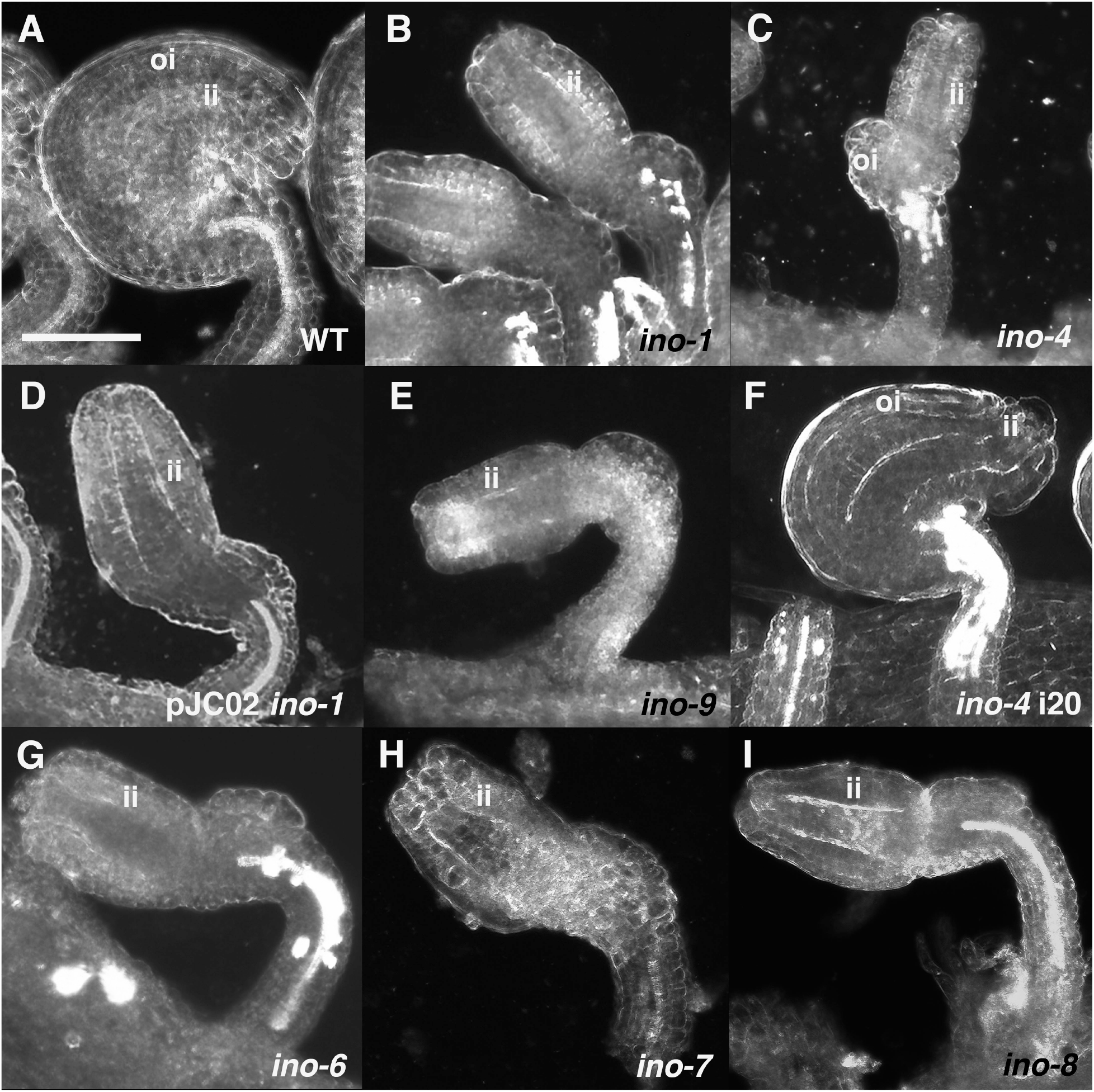
Dark field micrographs of Arabidopsis ovules. A) Wild-type. B) *ino-1*. C) *ino-4*. D) *ino-1* transformed with pJC02 transcribing *ino-4* mutant cDNA. E) *ino-9*. F) *ino-4* i20 double mutant. F) *ino-6*. H) *ino-7*. I) *ino-8*. oi, outer integument; ii, inner integument. Bar in A) is 50 µm in all panels. ii, inner integument; oi, outer integument.

### *ino-4* produces a small amount of wild-type *INO* mRNA

The finding that the *ino-4* aberrant mRNA/protein could not complement *ino-1* indicates there must be another source of active INO provided by the *ino-4* allele. Reverse transcriptase polymerase chain reaction (RT-PCR) analysis of mRNA from *ino-4* detected a prominent product of the expected size for the aberrant *ino-4* mRNA, but also a smaller amount of a product that co-migrated with the product derived from mRNA from wild-type plants (Fig. 3). To test if this product is properly spliced *INO* mRNA we devised primers that cross the splice junction and would amplify only the properly spliced *INO* cDNA (Supplemental Table S1). We found that *ino-4* does produce wild-type mRNA, along with the mis-spliced *ino-4*-specific product (Fig. 3).

**Fig. 3.**
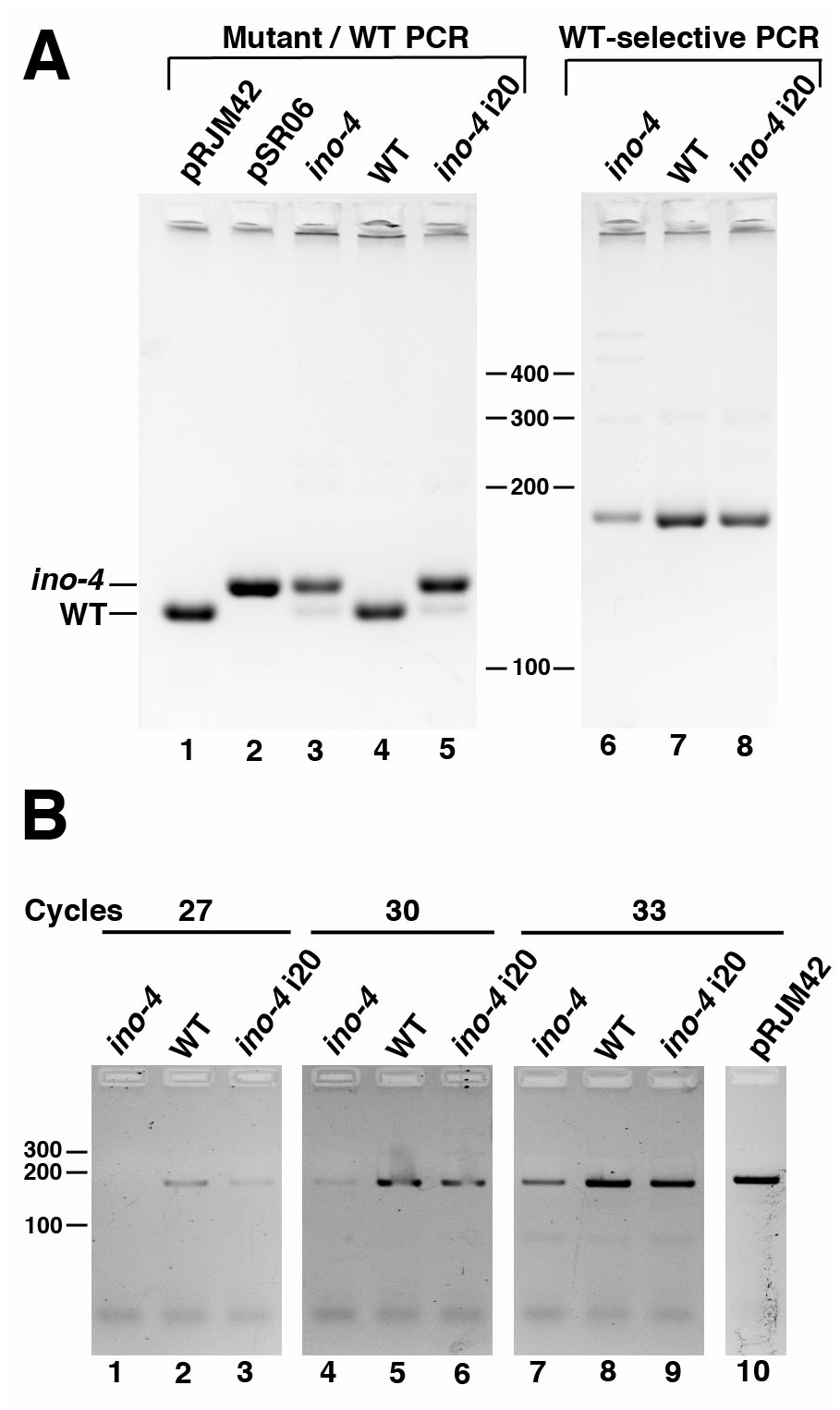
Reverse transcription PCR detection of *INO* mRNA. A) Lanes 1 – 5 show PCR products from primers flanking intron 4 (ino-4 FOR, ino-4 REV) that will amplify cDNA from both mRNA that utilized the wild-type splice site (WT), or the ectopic *ino-4* splice site (*ino-4*). Lanes 6 – 8 show products from primers that will selectively amplify only cDNA from mRNA using the wild-type splice site due to one primer crossing the junction between exons 4 and 5 (inoex45F2 with inoRTR1). Input DNA in lanes: 1) plasmid pRJM33 which includes the wild-type *INO* cDNA. 2) pSR06 which includes the cDNA to the aberrantly spliced *ino-4* cDNA. 3 and 6) cDNA to mRNA from *ino-4* flowers. 4 and 7) Wild-type Col-0 flower cDNA. 5 and 8) cDNA to mRNA from *ino-4* i20 double mutant flowers. B) PCR of cDNA from mRNA from flowers of the indicated plants or plasmid control (pRJM42) using primers selective for only the wild-type form of the *INO* mRNA. Samples were taken and evaluated following 27 cycles (lanes 1 – 3), 30 cycles (lanes 4 – 6), or 33 cycles (lanes 7 – 10). Sizes of DNA markers are indicated in base pairs at center (A) and left (B).

### *INO* quantitatively affects ovule development

The effect of the apparent reduced level of functional INO in *ino-4* suggests that the quantity of INO differentially affects outer integument growth. A test of this hypothesis was serendipitously produced by our identification of a second-site suppressor of *ino-4* mutants. *ino-4* plants produce small numbers of viable seeds which served as a starting point for suppressor screening. We mutagenized *ino-4* seeds with fast neutrons and screened M2 plants for increased outer integument growth and increased fertility. One homozygous *ino-4* line, i20, had significantly more outer integument growth than *ino-4* single mutants (Fig. 1F, 2F). We evaluated the presence of wild-type *INO* mRNA in *ino-4* and *ino-4*; i20. *ino-4*; i20 also produced both aberrant *ino-4* mRNA and wild-type mRNA (Fig. 3), but both were at an apparently higher level than in *ino-4* (Fig. 3).

Using segregating plants from a cross of *ino-4*; i20 (in the Col background) to L*er* we mapped the mutation in i20 to the top of chromosome 3. We used whole-genome sequencing to compare *ino-4*; i20 sequence to the TAIR10 Col reference sequence (Lamesch et al., 2012). We found an apparent deletion within the mapped region on the basis of the absence of sequence reads (Fig. 4A). This 57 bp deletion is between At3g10490 and At3g10500. Individual reads that spanned this region were not present, and examination of the sequence reads at the borders of the deletion on chromosome 3 showed that the reads paired with these reads were not aligned with chromosome 3. Instead reads from one side of the deletion paired with reads mapped to a region on chromosome 1, and those at the other end of the deletion paired with reads mapped to a region on chromosome 2. Examination of mapped reads in these regions of chromosomes 1 and 2 showed an approximately two-fold increase in depth of sequence reads, suggesting that these regions were duplicated in the genome. Two areas of chromosome 1 showed this duplication: a ∼ 40 kb region close to the *INO* locus, and a second ∼1.1 Mb region that included the *ino-4* sequence (Fig. 4B). Similarly, this section of chromosome 2 showed two-fold more reads mapped over a 2.2 kb region (Fig. 4C). The simplest explanation for this would be for regions of chromosomes 1 and 2 to be duplicated and inserted into a small deletion on chromosome 3. We used BLAST (Altschul et al., 1990) searches of our entire set of i20 reads to identify individual reads that crossed the borders between the deletion site and the putative insertions and were able to find the expected numbers of such reads. Applying this method recursively, we were able to determine that a small region of chromosome 2, and a larger portion of chromosome 1, that included the *ino-4* locus, were duplicated and inserted into chromosome 3 (Fig. 4D and Supplemental Fig. S1). Thus, the genome of i20 included an extra copy of the *ino-4* locus on chromosome 3. Using variable cycle counts in PCR we found that the level of wild-type mRNA was apparently higher in *ino-4*; i20 than in *ino-4*, but was still lower than in wild-type plants (Fig. 3B). Thus, an increase in the level of wild-type mRNA in *ino-4* i20 over the minimal level present in *ino-4* induced increased outer integument growth and produced ovules closer in form to those of wild-type plants. This demonstrates that INO activity role in ovule development is not a binary switch, but rather the activity quantitatively effects development of the outer integument.

**Fig. 4.**
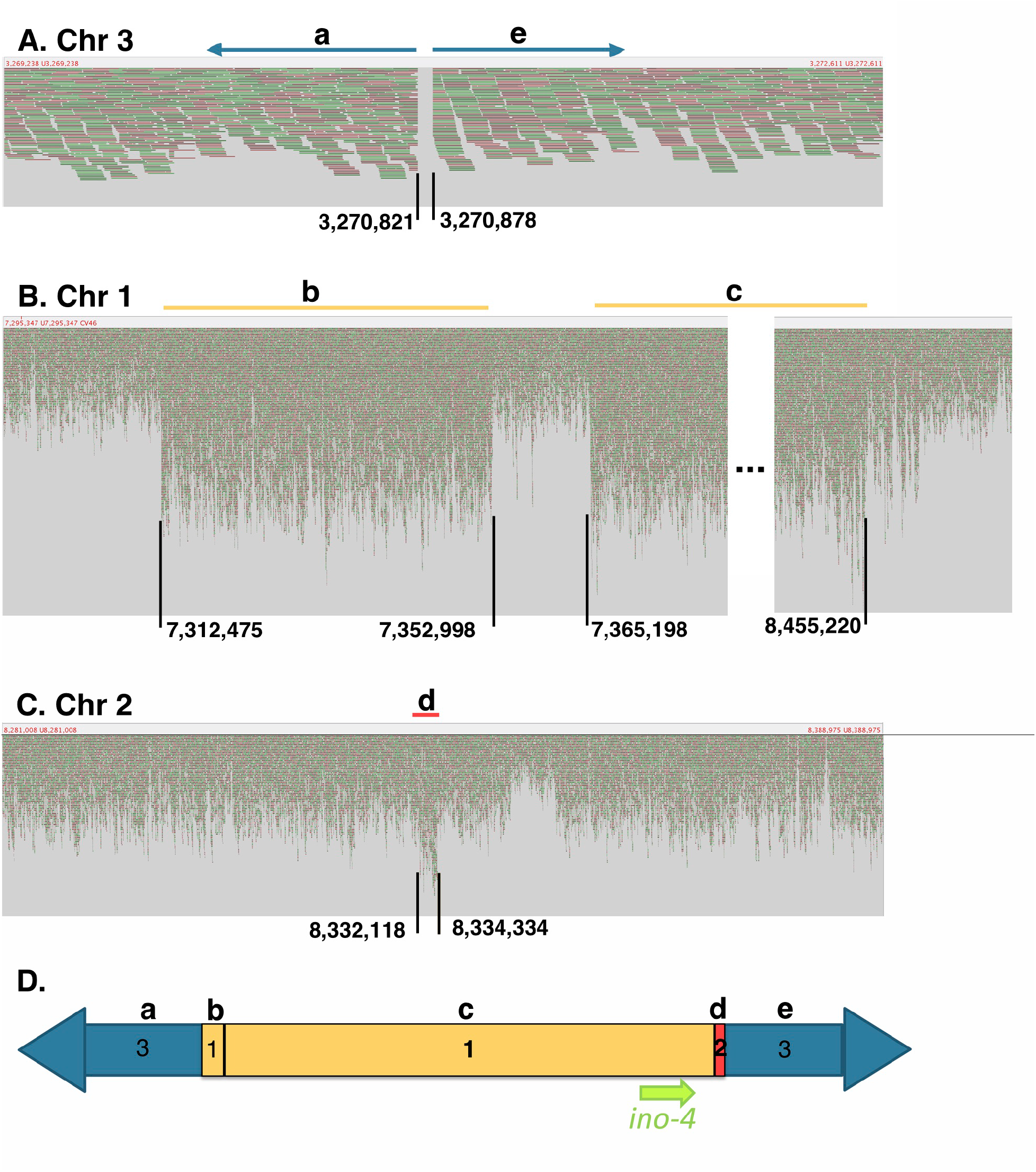
The structure of the insertion in the i20 region. A-C) Illumina sequence reads of homozygous *ino-4* i20 DNA aligned with wild-type Columbia Arabidopsis chromosomal regions as visualized in Tablet (Milne et al., 2013) software. Each ∼100 base sequence read is shown as a colored line (appearing more as colored spots at the zoomed-out view in B and C) aligned below the corresponding sequence of the indicated Arabidopsis chromosome. A) An absence of reads aligning with sequence between bases 3,270,821 and 3,270,878 of chromosome 3 indicates deletion of this region which is in the area within which i20 was mapped. Reads immediately to the left and right side of the deletion were paired with locations on chromosomes 1 and 2 respectively indicating the presence of inserted DNA. B) Two regions of chromosome 1 show approximately 2-fold more reads than expected. Reads at 7,312,475 paired with reads on chromosome 3 at position 3,270,821. Reads at position 8,455,220 paired with reads on chromosome 2 at position 8,332,118. C) A region of chromosome 2 also showed approximately 2-fold as many reads as an average region of the genome. Reads at 8,334,334 paired with reads on chromosome 3 at position 3,270,878. D) Diagram of the structure of the insertion into chromosome 3 of i20 that is composed of regions of chromosomes 1 and 2 (numbers indicate source chromosomes of the regions). Lower case letters at the top of the figure correspond to the chromosomal regions indicated by the same letters in panels A – C. The location of the *ino-4* gene in an inserted chromosome 1 region is indicated by the green arrow.

### Insertion mutant of *INO*

Since we had failed to isolate a clear null mutant of *INO* from chemically mutagenized material, we examined the set of available Arabidopsis insertion mutants for insertions in the *INO* locus. We found only one line, SALK_0063818, which we designated *ino-9*. Sequencing of genomic DNA flanking the inserted T-DNA shows that the insertion is 162 bp 5’ of the translation start site of the *INO* coding region. *ino-9* has a phenotype that precisely duplicates that of *ino-1* (Fig. 1, 2). The insertion is between the regulatory region of the promoter and the coding sequence (Simon et al., 2012), apparently interrupting expression. However, like the *ino-1* mutation, this mutation does not preclude the possibility of formation of active INO protein as initiation of transcription induced within the insertion could still produce wild-type INO protein. As a result, we still did not have certainty that the *ino* null mutant phenotype had yet been observed.

### CRISPR-induced null *ino* mutants

In the absence of identifying a verified null mutant from random mutagenesis we used CRISPR technology to induce deletion and insertion mutations in the coding region of *INO* that would cause frame-shifts and preclude formation of the full INO protein. A plant transformation plasmid, pGD60, was constructed to produce the CAS9 protein and two guide RNAs directed to two locations in the coding region of *INO*, and used to transform wild-type plants. The vector used included a cotyledon-expressed cassette producing green fluorescent protein (GFP). This enabled simple isolation of M2 seeds which had segregated out the CRISPR vector on the basis of the absence of GFP fluorescence (Aliaga-Franco et al., 2019). Multiple new *ino* mutant lines were isolated in the L*er* background, and three were chosen for complete characterization and designated *ino-6, -7*, and *-8. ino-6* included a single base deletion of a “T” residue at position 80 (in the first exon), and a second mutation inserting a “T” residue after position 883 (in exon 5) of the of the coding region of *INO*. This would introduce a frame shift and cause mistranslation of the entire zinc-finger region and the first half of the YABBY domain (Fig. S2). *ino-7* had an insertion of an “A” residue after position 883 (in exon 5) of the coding region, causing a frame shift of the second half of the YABBY domain (Villanueva et al., 1999) and all subsequent amino acids (Fig. S2). *ino-8* had a deletion of bases 77 and 78 of the coding region (in exon 1) of *INO*. This two-base deletion would cause a frame-shift leading to mistranslation of all of the INO protein excepting the first twenty-five amino acids (Fig. S2). All three of these mutants eliminate regions of INO shown to be critical to its function (Meister et al., 2005) and hence are null mutations. Homozygous plants of all three mutants were indistinguishable from *ino-1* plants and wild-type plants in vegetative habit, and all three produced ovules that duplicated the *ino-1* and *ino-9* phenotypes (Fig. 2). This result shows that *ino-1* and *ino-9* are also null mutants. Thus, the effects of *ino* complete loss of function mutants are confined to the outer integument of ovules, confirming the results of Vijayan et al. (2021).

## DISCUSSION

### *ino* null mutants

Until 2021, the two identified *ino* lesions had base substitutions that caused ectopic utilization of upstream splicing sites in *INO* pre-mRNA (Villanueva et al., 1999). Both retained the normal splice junctions and so had the potential for production of wild-type mRNA. This raised the possibility that null mutants had not been isolated. This failure to identify null mutants could result from such mutants being selected against due to pleiotropic effects. We first searched for null mutants in libraries of existing insertion mutants, but there found only a mutant with an insertion in the 5’-untranslated region. We could not say this was certainly null. We followed this by using CRISPR-based mutagenesis to produce frame-shift mutations in *INO* exons. Following completion of our mutagenesis, Vijayan et al. (2021) published work that included isolation of a putative null CRISPR-induced allele (*ino-5*). They reported the phenotype of *ino-*5 as not significantly different from *ino-1*. Confirming this result, all three of our CRISPR-induced null mutants, and also the insertion mutant, duplicated the *ino-1* phenotypic effects. This indicates that *ino-1* also was null.

The null mutants all exhibited complete failure in formation of an outer integument ((Villanueva et al., 1999; Vijayan et al., 2021); Fig. 1). All of the existing null mutants also exhibit only rare, sporadic formation of functional embryo sacs (Villanueva et al., 1999; Skinner and Gasser, 2009; Vijayan et al., 2021). In addition, the detailed imaging used by Vijan et al. (2021) revealed subtle alterations in formation of the nucellus and inner integument. However, expression of *INO* has only been observed in the abaxial layer of the outer integument and never in these other affected cells and locations (Villanueva et al., 1999; Balasubramanian and Schneitz, 2000). Vijan et al. (2021) hypothesize that absence of the outer integument would alter the hormonal environment during nucellar development. We note that allocation of nutritional resources would also be altered by absence of the outer integument, which would be expected to be a significant nutrient sink. Additionally, cell wall tension and other hydrostatic forces within the developing ovule would be altered by the absence of the confining outer integument and such forces also have roles in morphogenesis (Green, 1996). On this basis we concur with Vijan et al. (2021), that the effects of *ino* mutations on the development of the inner integument, nucellus, and embryo sac are most likely to be secondary effects of the absence of the outer integument, and not direct effects of absence of active INO protein.

It is interesting to note the only other plant in which an *ino* mutant has been identified to date was *Annona squamosa*, the sugar apple tree. In this *Thai seedless* (*Ts*) mutant the embryo sac still formed in the majority of ovules despite the complete absence of an outer integument (Lora et al., 2011). In Arabidopsis the tenuinucellate ovules have only a single nucellar cell layer surrounding the embryo sac, and this is sheathed in a thin two cell layer thick inner integument (Robinson-Beers et al., 1992; Vijayan et al., 2021). In contrast, the crassinucellate ovules of *A. squamosa* are surrounded by a robust nucellus of at least four cell layers and an inner integument comprising multiple cell layers at their micropylar end (Lora et al., 2011). Thus, even when the outer integument is absent there is substantial maternal tissue surrounding the developing embryo sac that could insulate it from the effects of the absence of an outer integument. This could explain the further development of the embryo sac in the *ino* mutant in this species relative to Arabidopsis. The *ino* mutation in *A. squamosa* appears to be a complete deletion of the *INO* locus, and hence is a null mutation (Lora et al., 2011). The absence of pleiotropic effects on other aspects of development in *A. squamosa* is thus consistent with the absence of pleiotropic effects now demonstrated for Arabidopsis *ino* null mutants.

### *INO* expression level corelates with the degree of outer integument growth

It had been previously hypothesized that the partial outer integument growth in *ino-4* resulted from a partially active protein produced from the mis-spliced, but in-frame *ino-4* mRNA (Villanueva et al., 1999). The failure of a transgene transcribing the cDNA of the mis-spliced mRNA to induce any detectable outer integument growth demonstrated that this was not the case. The mis-spliced mRNA produces a protein with a five amino acid insertion in the YABBY domain (Villanueva et al. 1999) and our result shows that this protein is not active, supporting the importance of the integrity of this domain for INO function. Instead, we were able to show that *ino-4* produces a small amount of properly spliced mRNA (Fig. 3). That this low amount of *INO* mRNA is able to promote a minimal growth of the outer integument (OI) is an indication that INO quantitively affects OI growth. Further evidence of this is provided by our isolation of a line with a translocated duplicate *ino-4* gene. As a result of the presence of this additional gene copy, this line has an increased level of wild-type mRNA over *ino-4* (Fig. 3) and an apparent proportional increase in OI growth (Fig. 1F, 2F). Thus, INO is a direct quantitative regulator of OI growth rather than being a binary switch. This is consistent with the function of *INO* in wild-type plants. *In situ* hybridization and reporter transgene studies (Villanueva et al., 1999; Meister et al., 2002) show that *INO* is expressed at the highest level on the gynobasal side of the ovule where maximal OI growth takes place. The level of *INO* expression shows a progressive decrease toward the gynoapical side of the ovule, correlating with the decreased OI growth in this area that leads to the hood-like outer integument of wild-type ovules (Meister et al., 2002). In *superman* (*sup*) mutant ovules there is increased OI growth on the gynoapical side of the ovule relative to wild-type (Gaiser et al., 1995). This correlates with an increase in *INO* expression in this region in such mutants (Meister et al., 2002). The correlation between developmental evidence of quantitative effects of *INO* expression and our mutant studies linking quantity of *INO* expression and amount of OI growth provides robust evidence that the level of *INO* expression is a primary determinant of OI growth.

## Conclusion

The phenotypes of the null *ino* mutants demonstrate that this gene’s role is specific to OI development in Arabidopsis. The specificity of the effects is mirrored in the confinement of *INO* expression to the outer (abaxial) layer of cells of the outer integument (Villanueva et al., 1999; Balasubramanian and Schneitz, 2002; Meister et al., 2002). This pattern of expression is conserved for *INO* orthologs among all examined angiosperms including other eudicots such as *Impatiens* and tomato (McAbee et al., 2005; Skinner et al., 2016), members of earlier-diverging groups such as magnoliids in the *Annona* genus (Lora et al., 2011) and the very earliest diverging angiosperms including *Cabomba* and *Amborella* (Yamada et al., 2003; Arnault et al., 2018). Functional analysis of *INO* activity has been evaluated through virus-induced gene silencing in tobacco (Skinner et al., 2016) and through analysis of an *INO* deletion mutant in *Annona* (Lora et al., 2011). Where effects similar to effects on Arabidopsis ovules were observed. Together these results indicate that *INO* has had a primary role as a regulator of OI development from the inception of the angiosperms. The OI is a synapomorphy of angiosperms and is one of the defining features separating angiosperms from gymnospermous progenitors (Gasser and Skinner, 2019; Shi et al., 2021). Phylogenetic analysis of the YABBY gene family shows that the origin of the *INO* clade correlates with the origin of the angiosperms (Yamada et al., 2011; Gasser and Skinner, 2019). Thus the origin of this critical or master gene for OI development (as shown by developmental and genetic analysis) may have been an essential step in the evolution of the OI and the origin of angiosperms (Gasser and Skinner, 2019; Shi et al., 2021).

## ACKNOWLEDGEMENTS

We thank Mily Ron and Anne Britt (Dept. of Plant Biology, U. C. Davis) for the gift of vectors and protocols for CRISPR mutagenesis, and Selina Rodrigues, Jennifer Zonka and George Day for excellent technical assistance. This work was supported in part by U. S. National Science Foundation grant IOS1354014 (to CSG).

## AUTHOR CONTRIBUTIONS

Work was planned and conceived by D. J. S. and C. S. G. D. J. S. performed fast neutron mutagenesis and analysis. C. S. G. and T. D. performed CRISPR mutagenesis and analysis. C. S. G. wrote the manuscript with editing by D. J. S. All authors approved the manuscript.

