## Supplemental Table and Figures for "The Arabidopsis *INNER NO OUTER (INO)* gene acts exclusively and quantitatively in regulation of ovule outer integument development"

**Supplemental Information**
Skinner et al., *INO* quantitatively regulates ovule development.

**Table S1** Oligonucleotides used

| **Name** | **Sequence** | **Purpose/Use** |
|---|---|---|
| AtINOcDNA-F | TGGATCCATGACAAAGCTCCCCAACATG | 5' primer for entire INO cDNA amplification, introduces BamHI site near start codon. |
| AtINOcDNA-r | TTCTAGATTACTCAAATGGAGATTTTCCCCA | 3' primer for entire INO cDNA amplification, introduces XbaI site following stop codon. |
| INOcds ex4/5F | AATTGCTTCATCAGGAAGAGATCAGG | Primer that crosses exon 4/5 boundary and will specifically amplify wild-type spliced sequence at this site. |
| INO RT R1 | TCTTCCTCACAAAAACATTGATCTG | Primer to pair with INOcds ex4/5F |
| ino4For | AAGCTTCCTTGTGAGCCATGC | Primer for amplification of region of INO cDNA including the aberrant *ino-4* splice site. Will amplify both wild-type and *ino-4* cDNA. |
| ino4Rev | TGAGAAGCGACAAAGAGCTCC | Primer for amplification of region of INO cDNA including the aberrant *ino-4* splice site. Will amplify both wild-type and *ino-4* cDNA. |
| InoCrisF4 | ATTGTAGTGGTGCAAAAACCACAC | Guide RNA targeting sequence one strand of INOCRIS4 |
| InoCrisR4 | AAACGTGTGGTTTTTGCACCACTA | Guide RNA targeting sequence other strand of INOCRIS4 |
| InoCrisF5 | ATTGAAAGGCTCAGAATCCAAGCA | Guide RNA targeting sequence one strand of INOCRIS5 |
| InoCrisR5 | AAACTGCTTGGATTCTGAGCCTTT | Guide RNA targeting sequence other strand of INOCRIS5 |

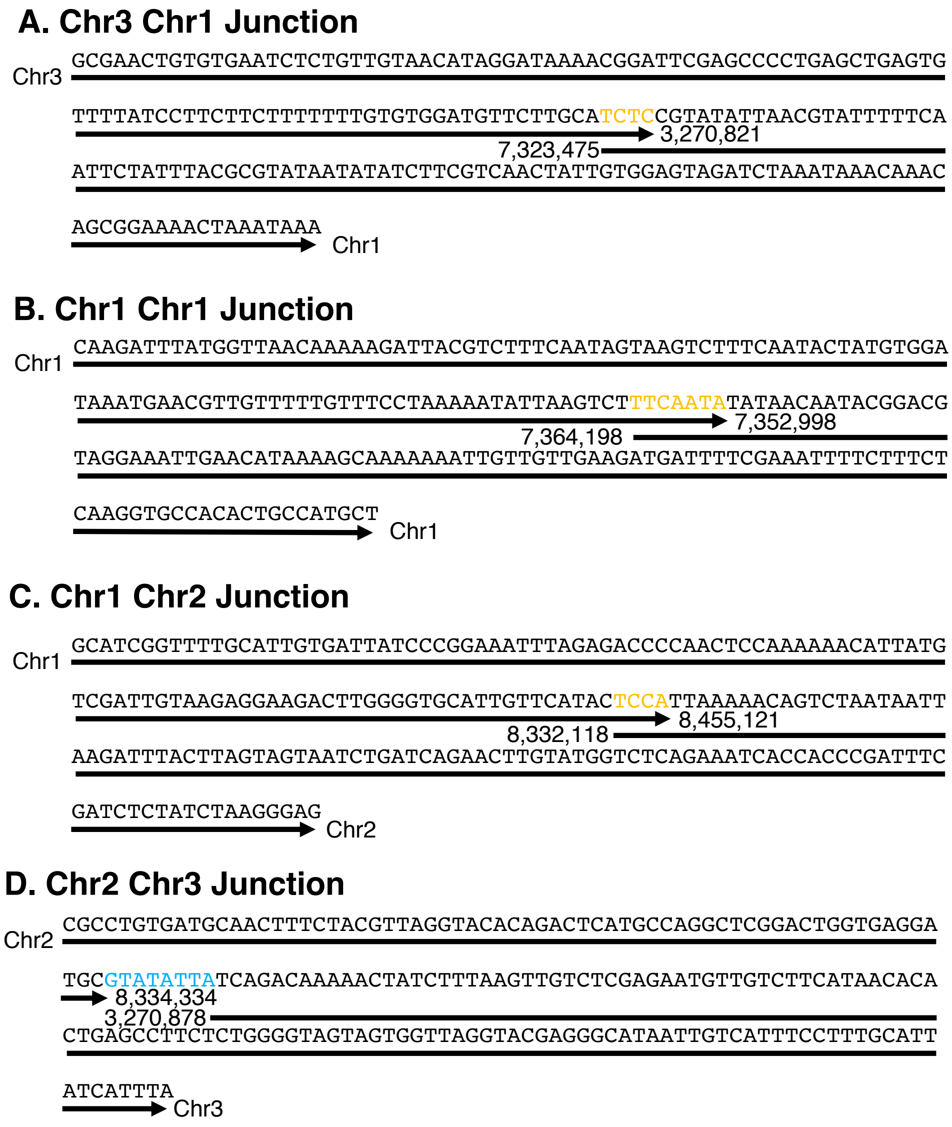


**Fig. S1**. Sequences of junction regions in the i20 translocated suppressor region. Sequences were assembled from Illumina sequence reads spanning and flanking the junctions. Regions spanning the junctions between (A) chromosomes 3 and 1, (B) chromosome 1 and 1, (C) chromosomes 1 and 2, and (D) chromosomes 2 and 3 as described in Fig. 4. In each panel the arrows under the sequences indicate the source chromosomes of that part of the sequence according to TAIR10 (Lamesch et al., 2012). The location of the junction ends of the sequences in those chromosomes in TAIR10 are indicated at the ends of the junction fragments. Orange bases are common to both chromosomes at the junction, and the aqua bases are an insertion of unknown origin at the junction.

```
gagactatgaagattccaattattacaagtgagatagagacgtacataagttgaattgtattgtgcattttagcata

          |(ino-9,T-DNA insert)
taaccaaagcattccacatgaaagatataaatataattacatatttgcattaatgaatattaacaaaacctatctct

                                                              START->
ctgtctatatatacacacacacacacacacatatatctcctatcattcatcgtacacacacactctctATGACAAAG

                                                              :(ino-6)
                                                             ::(ino-8)
CTCCCCAACATGACGACAACACTCAACCATCTATTTGATCTGCCGGGGCAGATTTGCCATGTCCAGTGTGGTTTTTG
CACCACTATTTTGCTGgtataccttaacttcttctcttattctctcccttcctttctcccttcctctctttatatat
atgatcatgagtttaatgagtgtaaatgtgttgcagGTGAGTGTACCGTTTACAAGCTTGTCAATGGTGGTGACTGT
GAGATGTGGGCATTGCACAAGCCTTCTCTCTGTCAATTTGATGAAGGCTTCCTTCATTCCTCTCCATCTCCTTGCTT
CTCTCTCCCATCTTGATGAGgtttccataacatctatacatagacaaacacacaccatctaaatatgaccaatatat
atatatatatacttacttttgattatatgtaaatatatgtgcagACCGGGAAAGAGGAGGTTGCAGCTACAGATGGT
GTGGAAGAAGAAGCATGGAAGGTGAATCAGGAGAAGGAGAACAGTCCAACGACTTTGGTTTCATCTTCAGACAATGA
AGATGAAGATGTGTCTCGTGTTTACCAAGTTGTCAATAAACgtacgtttgagtataaattcaacttaaactttagtt
ttgcgatttctcctttcatcagtcgtctaattatgttttgagtttgtatatatatagCACCTGAGAAGCGACAAAGA
GCTCCTTCAGCTTACAATTGCTTCATCAAgtaagaaataaattcaacacattacatgtgtatgtatattatatatat

                                                 a(ino-4 base change)
tcttacaaatttaaccacttaaggttttgagtttgtaaaatggttcttctttggggggttgtattggagcagGGAAGA

                         t(ino-6 base insert)
                         a(ino-7 base insert)
GATCAGGAGGTTAAAGGCTCAGAATCCAAGCATGGCTCACAAGGAAGCTTTCAGCTTAGCTGCCAAAAATgtactta
agcttaattagtcttaagatttaaaagaaaccctaatatagttttcatctttttgttagcttaattgacagatcga

                                   a(ino-1, ino-2 base change)
gttagctaataacattttatgatttcaatttttttgatttggggatggcgaagTGGGCCCATTTTCCTCCAGCTCACA
ACAAGAGAGCTGCTTCAGATCAATGTTTTTGTGAGGAAGATAACAATGCGATACTACCATGCAATGTTTTTGAGgta
atcctcagcaaccatttatattttttaatcactttcgcttcaaaatcaattaccatattcttgacaagtaaactcca
caacaagtgtcattttccaaagcaaaccaaataattaaagcattaactagatccaaatagcatcgctctcagtcaaa
atttcagaaccctagctagactaattatataaagagaaatcatttactaattattagagtgaaagcaaggagaaggg
taagtgagattgtgagaatatatcattgatcttcaactactacaatgtgtgaaatttatgacctaaacccttttata
gtattgaagcttacactgcacacagtttactttttatagaaacccattcaactaatttaaaacattgaagctgaatat
gtatagttttcaattaatataccaaataattgttgaaatgtttgtgatgcagGACCATGAAGAAAGCAATAATGGGT

                                                STOP
TCCGAGAGAGAAAGGCTCAGAGGCATTCCATTTGGGGAAAATCTCCATTTGAGTAAtaacaatttgggatatgaaaa
tttaacaaaaaaataaataatggcgtatgacatgtgatggtgactcttttcttctagggtttgattatgttggttta
gggtttctgttgttggagagagatagagagaagaagaatctgaaacggaagtaggatttgtgtgtgttcgagaagag
tccttgggaatgagcgagtttgttcagtaatttcgtagtacttgtcaactttgaacttaatttgaagagacatgcat
tgactctataaaaaaaattctacttttgcattcatgactactcttattccatgaatgttttaacattccatgaaaca
gtactttagtaagaatctttgttaacctatcataataaagatccttcttgacttcgcgcgctcaatg
```

**Fig. S2**. The Arabidopsis Col-0 *INO* gene sequence showing mutations. Regions are indicated by color and case. Untranscribed flanking regions are in lowercase black. 3’- and 5’-transcribed untranslated regions are in lowercase red. Introns are in lowercase lavender and protein coding sequences are in uppercase orange. Start and stop codons are highlighted in blue. Mutations *ino-1, -2,* and *-4* are from Villanueva et al. (1999) and other mutations are from this work. The mutations are indicated above the sequence with letters indicating substitution in place of the base they are above, or inserted following that base in the case of *ino-7* and the second mutation in *ino-6*. Colons indicate deletion of the bases they are above, and the vertical bar indicates the location of the T-DNA insertion in *ino-9.* All mutations are in the L*er* background excepting *ino-4* and *ino-9* which are in the Col-0 background.
